# Skeletal muscle effects of antisense oligonucleotides targeting glycogen synthase 1 in a mouse model of Pompe disease

**DOI:** 10.1101/2024.02.22.580414

**Authors:** Lan Weiss, Michele Carrer, Alyaa Shmara, Cheng Cheng, Hong Yin, Lac Ta, Victoria Boock, Yasamin Fazeli, Mindy Chang, Marvin Paguio, Jonathan Lee, Howard Yu, Angela Martin, Nina Raben, John Weiss, Tamar Grossman, Paymaan Jafar-nejad, Virginia Kimonis

## Abstract

Pompe disease (PD) is a progressive myopathy caused by the aberrant accumulation of glycogen in skeletal and cardiac muscle resulting from the deficiency of the enzyme acid alpha-glucosidase (GAA). Administration of recombinant human GAA as enzyme replacement therapy (ERT) works well in alleviating the cardiac manifestations of PD but loses sustained benefit in ameliorating the skeletal muscle pathology. The limited efficacy of ERT in skeletal muscle is partially attributable to its inability to curb the accumulation of new glycogen produced by the muscle enzyme glycogen synthase 1 (GYS1). Substrate reduction therapies aimed at knocking down GYS1 expression represent a promising avenue to improve Pompe myopathy. However, finding specific inhibitors for GYS1 is challenging given the presence of the highly homologous GYS2 in the liver. Antisense oligonucleotides (ASOs) are chemically modified oligomers that hybridize to their complementary target RNA to induce their degradation with exquisite specificity. In the present study, we show that ASO-mediated Gys1 knockdown in the Gaa^-/-^ mouse model of PD led to a robust reduction in glycogen accumulation in skeletal and cardiac muscle. In addition, combining Gys1 ASO with ERT further reduced glycogen content in muscle, eliminated autophagic buildup and lysosomal dysfunction, and improved motor function in Gaa^-/-^ mice. Our results provide a strong foundation for further validation of the use of Gys1 ASO, alone or in combination with ERT, as a therapy for PD. We propose that early administration of Gys1 ASO in combination with ERT may be the key to preventative treatment options in PD.

## INTRODUCTION

Glycogen storage disease type II, also called Pompe disease (PD) is an autosomal recessive disorder affecting between 1 in 17,000 and 1 in 40,000 individuals^1^. PD is caused by the excessive accumulation of glycogen in tissues resulting from inactivating mutations in the GAA gene, which encodes acid alpha-glucosidase, an enzyme responsible for the lysosomal degradation of glycogen. The pathological accumulation of glycogen affects many tissues, but it is by far most impactful in the heart and skeletal muscle of Pompe patients. Indeed, the lack of GAA enzyme activity, or extremely low levels (1–2% of normal), are associated with a fatal infantile cardiomyopathy, profound weakness, and skeletal muscle pathology, whereas 10-20% of normal GAA levels lead to skeletal myopathy with childhood or adult onset^2,3^. To understand the role of GAA in Pompe disease etiology, a mouse model has been generated by targeted deletion of the murine Gaa gene. The Gaa knockout (Gaa^-/-^) mouse recapitulates Pompe disease manifestations, including the accumulation of glycogen in skeletal and cardiac muscle, the autophagy defects, and the consequent myopathy^4^.

The standard of care for Pompe patients is enzyme replacement therapy (ERT) using recombinant human (rh) GAA (Lumizyme®, or Myozyme®, Sanofi Aventis US LCC), which became commercially available in 2006. ERT has shown positive results in improving the myopathy and cardiomyopathy associated with PD, and therefore has been used to help stabilize or partially improve patient outcomes^5–7^. However, ERT has limited efficacy in the skeletal muscle, as many patients continue to develop progressive muscle weakness despite receiving rhGAA enzyme^8–10^. Multiple factors contribute to the poor efficacy of ERT in the skeletal muscle of Pompe patients, particularly when significant disease pathology is present. Some of the limitations affecting ERT are associated with the complexity of delivering a relatively large protein to the muscle tissue.

Indeed, when administered via intravenous infusion, the recombinant GAA enzyme circulates to the liver, which acts as a major “sink” and precludes more than 80% of the administered protein from reaching the skeletal muscle^11^. Furthermore, ERT relies on the cation-independent mannose-6-phosphate receptor (CI-MPR), which is expressed at low levels on the membrane of skeletal muscle cells^7^. In addition, dysregulation of autophagy, and the consequent impairment of the autophagosome-lysosome fusion in Pompe myofibers, further contribute to the reduced efficacy of ERT. Ultimately, the failure of autophagy leads to inefficient trafficking of the therapeutic enzyme to the lysosome, limiting its ability to clear the glycogen^12^. This phenomenon is especially evident in type II muscle fibers, which are particularly refractory to ERT^10^. Previous studies performed in mice found upregulated protein levels of enzymes involved in glucose uptake and cytoplasmic glycogen synthesis in skeletal muscle from mice with Pompe disease, including glycogenin (GYG1), glycogen synthase (GYS1), glucose transporter 4 (GLUT4), glycogen branching enzyme 1 (GBE1), and UDP-glucose pyrophosphorylase (UGP2) suggesting a positive feedforward loop for cellular glycogen accumulation^13^. Administration of rhGAA to the Pompe mice normalized not only muscle glycogen levels but also G6P, hexokinase, GS, and glycogenin levels^14^. Thus, it would appear that correction of one aspect of the glycogen metabolic pathway (lysosomal glycogen) results in the correction of multiple other components. Studies in patient biopsies however found that extralysosomal glycogen accumulation in muscle biopsies from late-onset patients was not cleared by ERT, indicating the importance of an acidic pH environment^15^.

Glycogen synthase (GYS) enzymes catalyze the addition of glucose residues to the growing glycogen molecules by formation of α-1,4-glucosidic linkages. Two paralogous GYS genes, GYS1 and GYS2, exist in mice and humans. GYS2 expression is restricted to the liver, where it plays an essential role in the maintenance of proper body glucose homeostasis^16^. GYS1 has a broader expression profile, but it is particularly abundant in skeletal muscle and heart^16^. Decreasing GYS1 expression represents a potential therapeutic approach for PD, based on the principle of substrate reduction. Indeed, limiting the synthesis of new glycogen would result in attenuated glycogen accumulation in the muscle tissue. Genetic ablation of murine Gys1 provided proof of principle validation for substrate reduction therapy (SRT) in vivo. Knocking down Gys1 led to restoration of muscle functionality in the Gaa^-/-^ mouse model^17^. Similarly, intramuscular injection of recombinant adeno-associated virus-1 (AAV-1) vector expressing short hairpin ribonucleic acid (shRNA) targeted to Gys1 has been shown to reduce glycogen accumulation in the gastrocnemius of newborn Gaa^-/-^ mice ^18^. Altogether the preclinical evidence suggests that reducing GYS1 can be a potential therapeutic approach in Pompe disease, either alone or in combination with ERT. However, GYS1 and GYS2 show high degree of homology, which poses a complex challenge in the identification of safe, isoform-specific inhibitors^19^. It is important to note that no approved small molecule inhibitor is clinically available for the treatment of Pompe disease.

Antisense oligonucleotides (ASOs) are short synthetic nucleic acids that hybridize to complementary target RNAs in a sequence-specific manner using Watson-Crick base pairing. By doing so, ASOs can influence RNA processing and regulate protein expression via multiple mechanisms, including RNase H1-mediated mRNA degradation, and splicing modulation^20–24^. Over the past two decades ASOs have emerged as both powerful research tools and therapeutic molecules^25–29^. The intrinsic properties of ASOs make them the ideal pharmacological tool to reduce muscle GYS1 expression with exquisite specificity.

Here, we show that systemic treatment with a phosphorothioate (PS) 2’-constrained ethyl (cEt)-modified gapmer ASO targeting murine Gys1 not only reversed the excessive glycogen accumulation, but also corrected the autophagic defects present in the skeletal muscle of Gaa^-/-^ mice. The ASO-mediated reduction of GYS1 protein, especially when combined with ERT, led to restoration of muscle function in Pompe mice, thus underscoring the in vivo potential of our approach.

## RESULTS

### Progressive glycogen accumulation and muscle weakness in the Gaa^-/-^ mouse model

We utilized the Gaa knockout mouse model to study the effects of lowering Gys1 expression on Pompe myopathy in vivo. The Gaa^-/-^ mouse was generated and studied by Raben et al. (1998)^4^. To inform the design of appropriate prevention and reversal treatment paradigms, we characterized the progressive accumulation of glycogen in the skeletal and cardiac muscle of the Gaa^-/-^ mice from our colony. We utilized two methods to measure glycogen content in the tissues: a colorimetric biochemical assay, and periodic acid-Schiff (PAS) histochemical staining^4^. As expected, we observed higher amounts of glycogen in the skeletal muscle of Gaa^-/-^ mice compared to wild type mice (Figure 1A). Remarkably, the glycogen levels in quadriceps muscle were already elevated in one-month-old Gaa^-/-^ mice, suggesting that the mutant mice start to accumulate glycogen in skeletal muscle at an early age.

**Figure 1.**
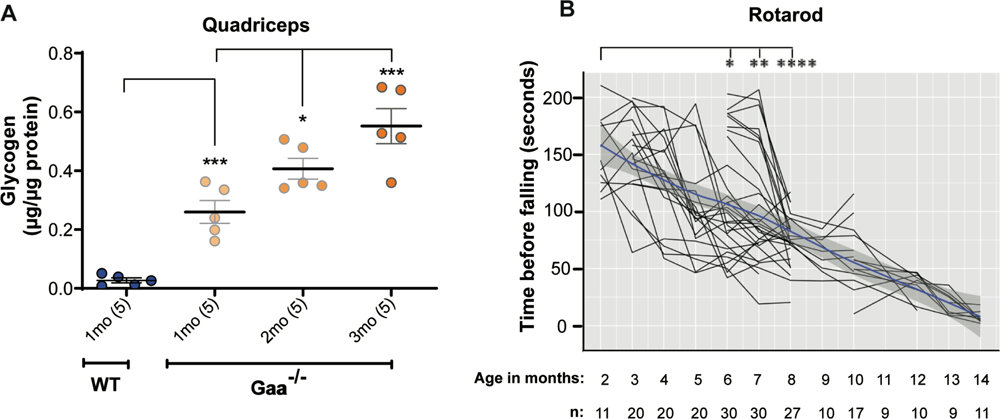
Gaa^-/-^ mice show progressive accumulation of glycogen in skeletal muscle followed by deterioration of muscle function. **A.** Glycogen content in quadriceps from Gaa^-/-^ mice at 1, 2, and 3 months (mo) of age compared to 1-month-old wild type (WT) C57BL/6 mice. n=5 per group, as indicated in parenthesis on the x-axis. Statistical analysis was performed using unpaired t-test in GraphPad Prism software. **B.** Graph illustrates the time that the mice spent on an accelerating rotarod before falling, to assess muscle strength as the animals age from 2 to 14 months. The mean decline in motor function is represented by the blue line within the shaded grey area, which indicates the SEM range. The number of mice in each measurement group is indicated at the bottom of the graph (n). Statistical analysis was performed using one-way ANOVA in GraphPad Prism software. *: p<0.05, **: p < 0.01, ***: p < 0.005, ****: p < 0.001.

The accumulation of glycogen increased progressively over time, from 0.26, to 0.41, and 0.55 µg glycogen per µg of total protein in quadriceps muscle homogenate from one-, two-, and three-month-old Gaa^-/-^ mice, respectively (Figure 1A). Similar to previous reports ^4,30,31^, the excessive glycogen accumulation in the skeletal muscle was associated with a progressive decline in motor function in aging Gaa^-/-^ mice, as assessed by the rotarod test (Figure 1B).

### Different levels of ASO-mediated Gys1 knockdown lead to the proportional amelioration of **aberrant glycogen accumulation in the skeletal muscle of young Gaa^-/-^ mice**

Through in vitro and in vivo screening, we identified two ASOs targeting mouse Gys1 that were potent and well tolerated in mice (Supplementary Figure 1, 2). When dosed systemically to wild type mice via subcutaneous injection for six weeks, Gys1 ASO#1 and ASO#2 resulted in the efficient, dose-dependent knockdown of murine Gys1 mRNA. ASO#1 had an ED_50_ of 45.2 mg/kg (95% confidence interval (C.I.): 32.3-63.2 mg/kg) in quadriceps muscle, whereas ASO#2 was more potent, with an ED_50_ of 24.2 mg/kg (95% C.I.: 22.4-26.1 mg/kg) (Supplementary Figure 1A). Consistently, Gys1 ASO#2 achieved a robust reduction of GYS1 protein in skeletal muscle of wild type and Gaa^-/-^ mice (Supplementary Figure 1B, 1C).

To determine if ASO-mediated Gys1 knockdown attenuates the pathological accumulation of glycogen in vivo, we dosed young Gaa^-/-^ mice with the newly identified Gys1 ASO#1 and ASO#2. A starting age of one month was selected because at this time the Gaa^-/-^ mice have not developed severe myopathy, yet, and the glycogen is only starting to accumulate in the skeletal muscle (Figure 1A). The one-month-old Gaa^-/-^ mice (Cohort 1) were dosed via subcutaneous injection with 25 mg/kg Gys1 or control ASO, once a week for sixteen weeks (Figure 2A). Compared to control ASO, Gys1 ASO#1 and ASO#2 resulted in 83.64% (95% C.I.: 77.40-88.58) and 88.58% (95% C.I.: 83.61-92.06) Gys1 mRNA knockdown, respectively in quadriceps muscle (Figure 2B, Supplementary Table 1). Corroborating the reduction in mRNA expression, compared to control ASO, ASO#2 administration decreased GYS1 protein, measured by Western blot, in key Pompe tissues of Gaa^-/-^ mice by 100%, 98%, and 100% in quadriceps, diaphragm, and heart, respectively. (Figure 2C, Supplementary Table 1).

**Figure 2.**
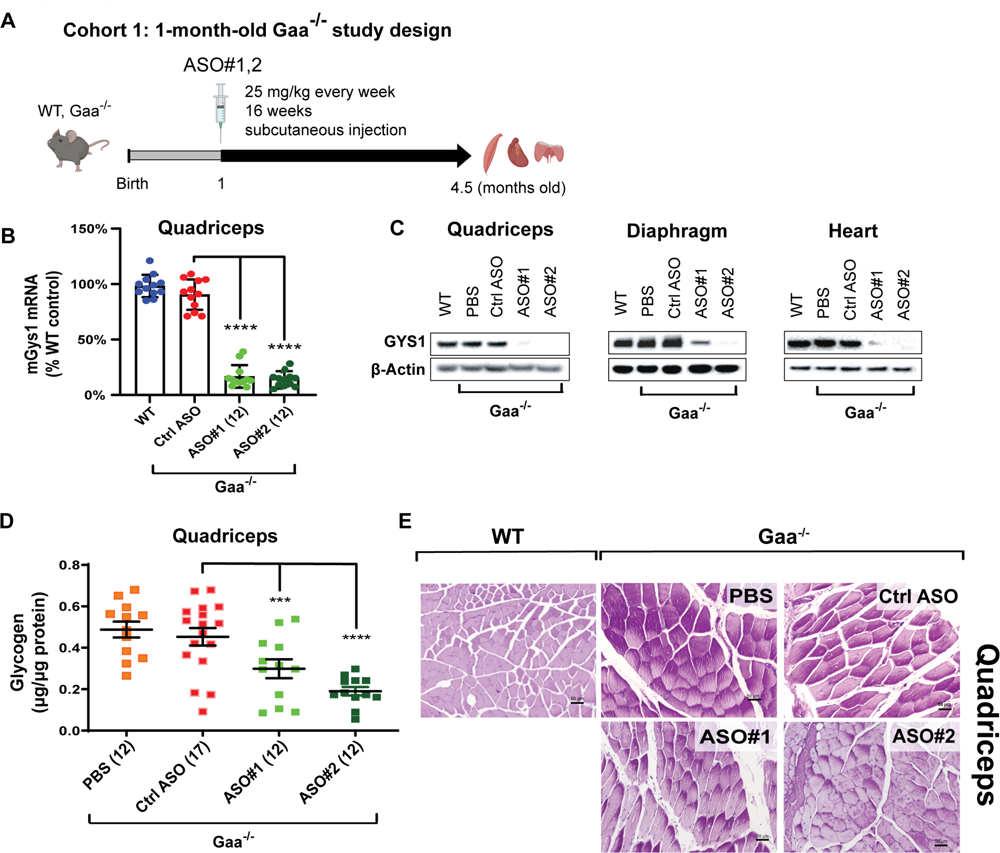
Early treatment of Gaa^-/-^ mice with Gys1 ASOs lowers target mRNA and protein expression, leading to reduced glycogen accumulation in skeletal muscle. **A.** Schematic illustration of the early treatment paradigm with Gys1 ASOs in Gaa^-/-^ mice from 1 to 4.5 months of age. WT: wild type. **B.** qPCR analysis of Gys1 mRNA in quadriceps muscle. **C.** Representative images of Western blot analysis of GYS1 protein in quadriceps, diaphragm, and heart. The number of samples in each group is listed in parentheses on the x-axis. **D.** Glycogen content measured using a biochemical assay in quadriceps muscle from Gaa^-/-^ mice dosed with either PBS, control ASO (Ctrl ASO), or two different Gys1 ASOs. The number of samples in each group is listed in parentheses on the x-axis. **E.** Periodic acid-Schiff (PAS) staining of histological sections of quadriceps muscle. Scale bars: 50 µm. Statistical analysis was performed using unpaired t-test in GraphPad Prism software. ***: p < 0.005, ****: p < 0.0001.

To evaluate the impact of Gys1 knockdown on excessive glycogen accumulation in the muscle of the Pompe mice, we measured glycogen content by a colorimetric biochemical assay in the muscle of the 20-week-old Gaa^-/-^ mice following 16 weeks of Gys1 ASO treatment. ASO#1 resulted in a 38.06% reduction in glycogen content (0.29 μg/μg protein) in quadriceps muscle, and ASO#2 resulted in 57.05% reduction in glycogen (0.19 μg/μg protein) compared to control ASO (0.49 μg/μg protein) (Figure 2D, Supplementary Table 1). The superior efficacy of ASO#2 compared to ASO#1 is consistent with its greater potency in knocking down Gys1 mRNA expression. Similarly, the control ASO had no effect on Gys1 expression and glycogen content (Figure 2B-E). In agreement with the biochemical measurements, semi-quantitative analysis of glycogen content in histological sections using PAS staining confirmed that Gys1 ASOs ameliorated the aberrant accumulation of glycogen in Gaa^-/-^ quadriceps muscle. On a scale of 1 (no staining) to 4 (extensive staining), an average score of 1.8 was assigned to the 4.5-month-old Gaa^-/-^ mice treated with control ASO, 1.4 to mice treated with ASO#1, and 1 to mice treated with ASO#2 (Figure 2E, Supplementary Table 2).

In view of the enhanced efficacy in the early-treatment setting, we selected ASO#2 to further investigate if Gys1 ASO treatment could revert the Pompe pathology in older Gaa^-/-^ mice.

### ASO-mediated Gys1 reduction partially reverses glycogen accumulation in skeletal muscle of older Gaa^-/-^ mice

To assess if therapeutic treatment with Gys1 ASO could reverse the glycogen accumulation and myopathy in mature Pompe mice, we dosed older Gaa^-/-^ mice with Gys1 ASO#2 starting at three months of age (Cohort 2) (Figure 3A). Gys1 ASO#2 was administered by weekly subcutaneous injections at 25 mg/kg for 16 weeks (Figure 3A). The rotarod motor performance assay showed that the ASO#2-mediated knockdown of Gys1 expression resulted in a trend towards preservation of motor function (Figure 3B), which otherwise progressively declines in Gaa^-/-^ mice (Figure 1B, 3B). Indeed, the control ASO did not slow down the deterioration of muscle function in Pompe mice (Figure 3B). We sacrificed the animals at 6.5 months of age and studied the quadriceps muscle, diaphragm, and heart tissue from mice treated with Gys1 ASO#2. We measured a 64.71% (95% C.I.: 57.00-71.05) knockdown of mouse Gys1 mRNA and complete depletion of GYS1 protein in quadriceps muscle compared to control ASO (Figure 3C-D, Supplementary Table 1). Similarly, GYS1 protein was reduced by 80% in diaphragm and 70% in heart compared to mice treated with control ASO (Figure 3D, Supplementary Table 1). Of note, the expression of GYS1 was not significantly different in 6.5-month-old Pompe mice compared to wild type mice (Figure 3D). A biochemical assay demonstrated that the ASO#2-mediated knockdown of Gys1 expression was associated with a modest 7.79% (95% C.I.: 5.53-28.45) reduction in glycogen accumulation compared to control ASO in the quadriceps muscle of Gaa^-/-^ mice (Figure 3E, Supplementary Table 1). In addition, semi-quantitative analysis of histological sections of quadriceps muscle using PAS staining confirmed a mild improvement of the aberrant glycogen accumulation following treatment with Gys1 ASO#2 (average score of 1.25 on a scale of 1 to 4) compared to control ASO (average score of 2.2) (Figure 3F, Supplementary Table 2).

**Figure 3.**
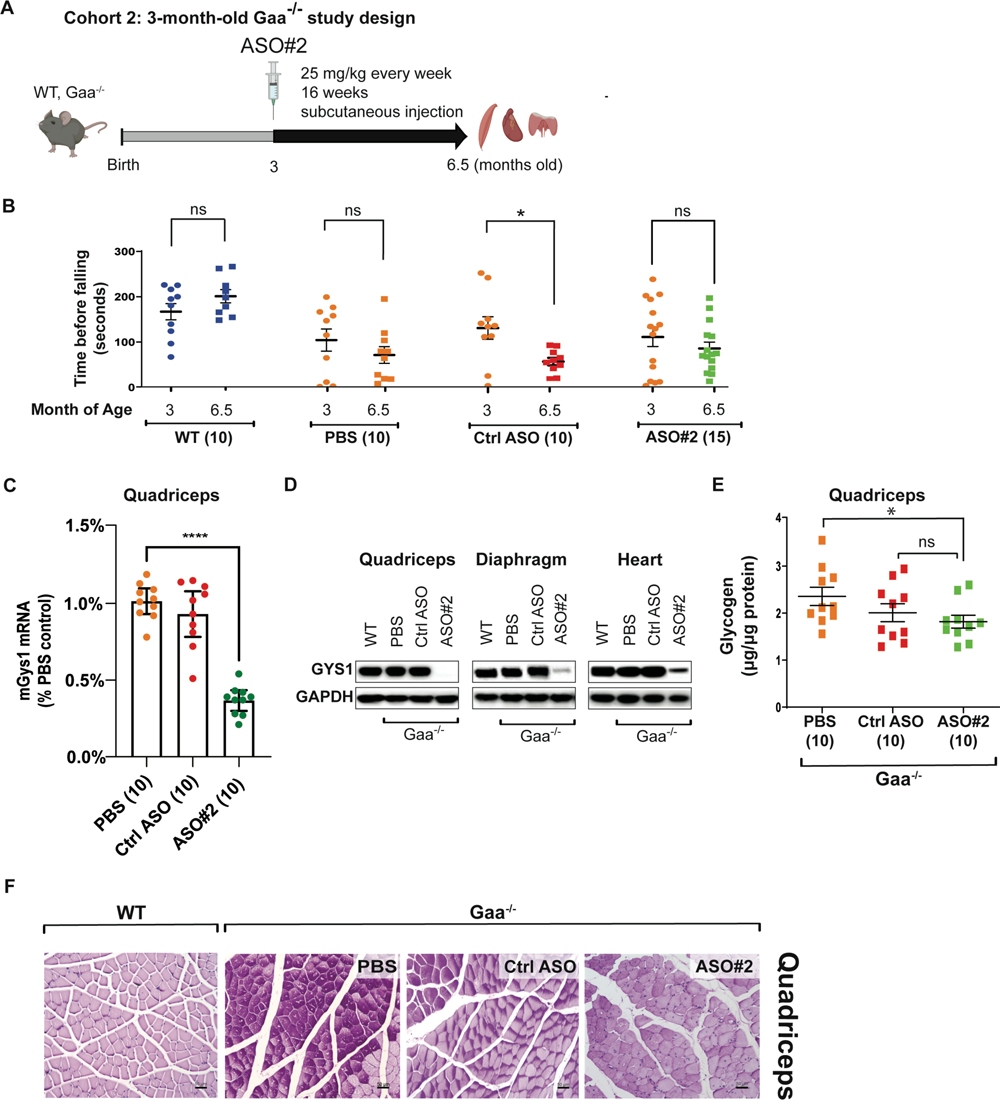
Therapeutic treatment with Gys1 ASO in Gaa^-/-^ mice partially reverses glycogen accumulation in skeletal muscle and preserves muscle strength. **A.** Schematic illustration of the reversal treatment paradigm with Gys1 ASO#2 in Gaa^-/-^ mice from 3 to 6.5 months of age. WT: wild type. **B.** Graph illustrates the time that mice spent on an accelerating rotarod before falling, to assess muscle strength. **C.** qPCR analysis of Gys1 mRNA. The number of samples in each group is listed in parentheses on the x-axis. **D.** Representative images of Western blot analysis of GYS1 protein in quadriceps, diaphragm, and heart. GAPDH was used as loading control. **E.** Glycogen content measured using a biochemical assay in quadriceps muscle from Gaa^-/-^ mice dosed with either PBS, a control ASO (Ctrl ASO), or Gys1 ASO. The number of samples in each group is listed in parentheses on the x-axis. **F.** Representative images from PAS staining of histological sections of quadriceps muscle. Scale bars: 50 µm. Statistical analysis was performed using unpaired t-test in GraphPad Prism software. *: p<0.05, ***: p < 0.005, ****: p < 0.001, ns: not significant.

### ASO mediated Gys1 knockdown in combination with enzyme replacement therapy further reverses Pompe pathology, preserving motor function in older Gaa^-/-^ mice

Substrate reduction therapy (SRT) using ASO to remove the source of new glycogen synthetized via GYS1 shows promise as a monotherapy, but the reversal of the myopathy in Gaa^-/-^ mice was incomplete in 6.5-month-old animals. We reasoned that combining Gys1 ASO, which blocks the synthesis of new glycogen, with GAA enzyme replacement therapy (ERT) to simultaneously remove pre-existing glycogen in the skeletal muscle and heart of older Pompe mice, could provide additive efficacy. Therefore, we dosed four-month-old Gaa^-/-^ mice with Gys1 ASO#2 at 25 mg/kg by weekly subcutaneous injections for six weeks, followed by a 10-week-long co-administration phase where we combined ASO#2 and ERT, in the form of recombinant human alpha-acid glucosidase (rhGAA) enzyme (Cohort 3) (Figure 4A). rhGAA was dosed intravenously every two weeks (Figure 4A). Alternatively, Gaa^-/-^ mice were dosed with either ASO#2 alone or ERT alone (Figure 4A). Wild type mice and PBS-treated Gaa^-/-^ mice were included in the study as controls. The ASO#2+ERT combination treatment led to a stable increase in rotarod performance over the 4-month-long treatment, resulting in gains of 9, 8, 11 and 19% compared to baseline after 1, 2, 3, and 4 months of treatment, respectively. On the other hand, ASO#2 alone increased the rotarod performance in the first three months of treatment, but was followed by a slowdown (2, 10, 12 and 7% gain compared to baseline after 1, 2, 3, and 4 months of treatment, respectively). ERT alone was unable to prevent the functional decline in the aging Pompe mice, resulting in a downward change in rotarod performance over time (-2, -8, -5 and -8% compared to baseline after 1, 2, 3, and 4 months of treatment, respectively) (Figure 4B).

**Figure 4.**
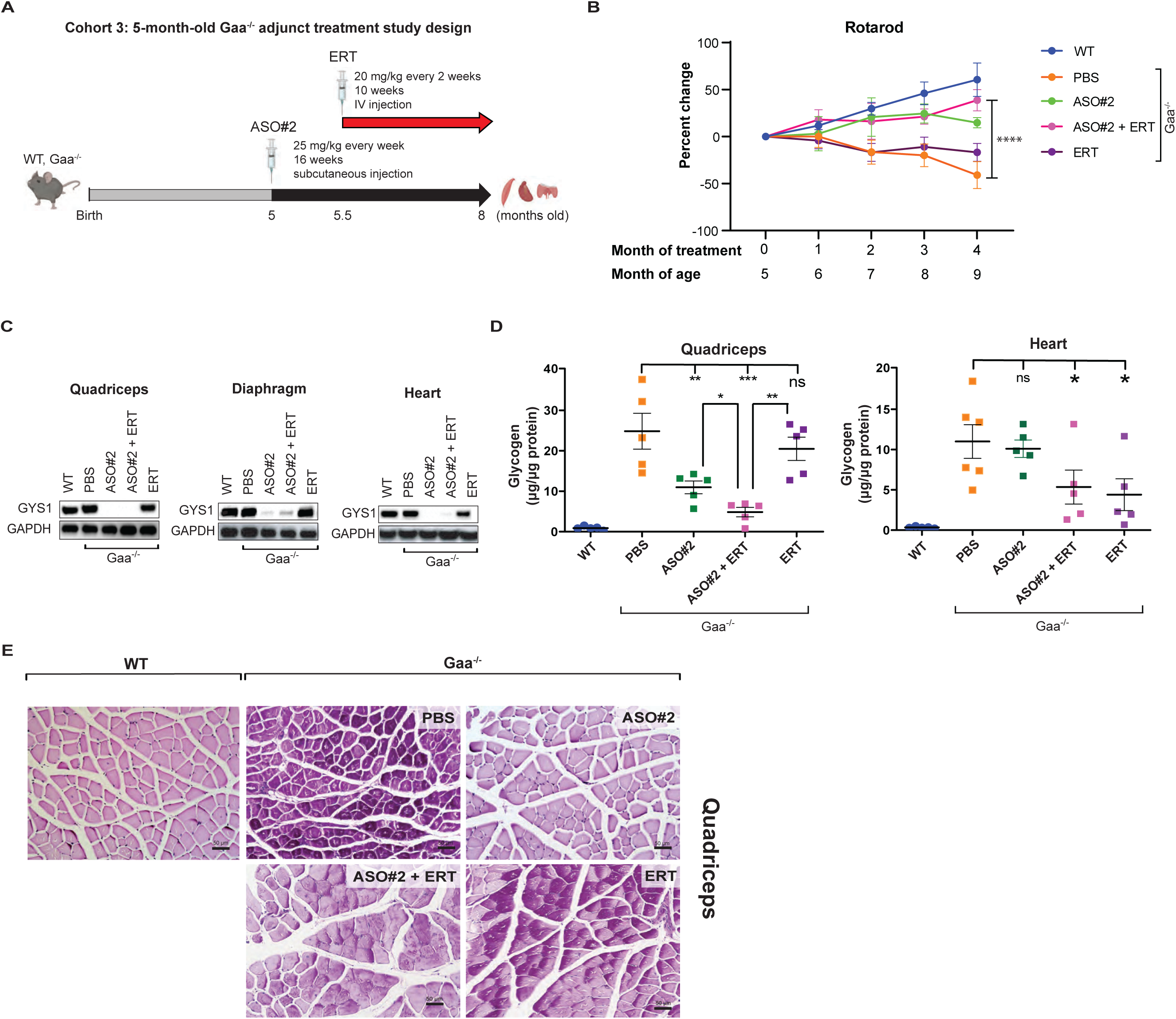
Combination treatment with Gys1 ASO and ERT reduces glycogen accumulation and improves motor function in Gaa^-/-^ mice. **A.** Schematic illustration of the study where 4-month-old Gaa^-/-^ mice were dosed with Gys1 ASO#2 and ERT in combination. WT: wild type. **B**. Graph shows the results of a rotarod test to assess muscle strength. The time that the mice spent on a rotating rod was recorded monthly over the three months of treatment and expressed as percent change versus baseline measurements (collected before the start of dosing) for each group; n=5 per group. Statistical analysis in B was performed by two-way ANOVA using multiple comparisons in GraphPad Prism software. ****: p < 0.001 between the Gaa^-/-^ mice that received PBS and Gys1 ASO#2+ERT. **C.** Representative images of Western blot analysis of GYS1 protein in quadriceps muscle, diaphragm, and heart. GAPDH was used as loading control. **D.** Glycogen content measured using a biochemical assay in quadriceps muscle and heart from wild type or Gaa^-/-^ mice dosed with either PBS, Gys1 ASO#2, Gys1 ASO#2+ERT, or ERT alone (n=5 per group). Statistical analysis in C and D was performed using unpaired t-test. *: p<0.05, **: p < 0.01, ***: p < 0.005, ns: not significant.

At the end of the study, the 8-month-old mice were sacrificed for further biochemical and IHC analysis of the muscle tissue. Western blot analysis revealed that ASO#2 alone or ASO#2+ERT resulted in the complete ablation of GYS1 protein expression in quadriceps muscle and heart (Figure 4C). Similarly, minimal residual GYS1 protein was present in the diaphragm of the same treatment groups (Figure 4C). On the other hand, Gaa^-/-^ mice that received PBS or ERT alone had robust GYS1 protein expression, comparable to wild type levels (Figure 4C). Importantly, the systemic administration of Gys1 ASO#2, alone or in combination with ERT, was well tolerated in Pompe mice with no significant changes in body or organ weights, and no significant elevation in liver transaminases, blood urea nitrogen, or muscle creatine kinase in plasma compared to control groups (Supplementary Figure 2).

The average glycogen content measured using the biochemical assay in quadriceps muscle of Gaa^-/-^ mice was 24.85 (95% C.I.: 12.54-37.17), 10.98 (95% C.I.: 6.57-15.38), 4.84 (95% C.I. 1.61-8.07), and 20.5 (95% C.I.: 12.46-28.53) μg glycogen per μg total protein in the PBS, ASO alone, ASO+ERT, and ERT alone treatment groups, respectively (Figure 4D). Therefore, compared to PBS controls, the biochemical glycogen assay revealed a 55.22% (95% C.I.: 20.29-74.84) reduction in glycogen content in quadriceps muscle of Gaa^-/-^ mice dosed with ASO#2 alone (Figure 4D, Supplementary Table 1). The ASO#2+ERT combination resulted in an even greater effect, reducing glycogen levels by 83.48% (95% C.I.: 53.80-94.10) in quadriceps muscle (Figure 4D, Supplementary Table 1). ERT alone did not significantly attenuate the glycogen accumulation, leading to 15.78% (95% C.I.: 11.61-31.49) less glycogen in quadriceps muscle of Gaa^-/-^ mice. ERT was more efficacious in the heart, where it resulted in 65.82% (95% C.I.: 5.88-87.59) glycogen reduction vs. PBS control when dosed alone, and in 74.93% (95% C.I.: 18.77-92.26) glycogen reduction vs. PBS control when combined with Gys1 ASO#2 (Figure 4D, Supplementary Table 1). PAS staining of quadriceps muscle histological sections corroborated the biochemical assay results, showing a marked reduction in glycogen content, especially in the ASO#2 group (semi-quantitative score of 1.5 on a scale of 1 to 4, where 1 is no glycogen detected) and ASO#2+ERT (score of 1), compared to PBS (score of 3.75) or ERT alone (score of 3) samples (Figure 4E, Supplementary Table 2). Thus, the motor function improvement correlated well with the reduction in muscle glycogen content.

Taken together, our data supports the hypothesis that blocking the synthesis of new glycogen while concomitantly enhancing its clearance is beneficial in treating both the skeletal muscle and cardiac abnormalities in Pompe disease (Figure 6).

### ASO-mediated Gys1 knockdown with or without ERT, but not ERT alone, ameliorates the autophagic buildup in the skeletal muscle of Gaa^-/-^ mice

Defective autophagy is a major factor contributing to the muscle pathology in Pompe disease^7^. Consistent with the highly integrated functions of lysosomes and autophagosomes, it has been shown that lysosomal dysfunction in Pompe disease leads to a block in autophagic flux and accumulation of autophagic debris, particularly in muscle tissue^32^. It is important to note that the Gaa^-/-^ mouse model faithfully recapitulates the autophagic defects described in Pompe patients^30^. Incomplete autophagic flux can be assessed by measuring the aberrant accumulation of LC3 and p62 proteins. Additionally, immunostaining of histological sections for lysosomal-associated membrane protein 1 (LAMP1) provides a method to detect enlarged lysosomes^33,34^. We investigated the efficacy of Gys1 ASO, alone or in combination with ERT (Cohort 3, Figure 4A), in ameliorating the defective autophagy in Gaa^-/-^ mice. Western blot analysis confirmed accumulation of p62 and LC3-II proteins in quadriceps muscle and heart of 8-month-old Gaa^-/-^ mice treated with PBS (Figure 5A). ERT alone alleviated the autophagic defect in the heart but not significantly in the skeletal muscle, as indicated by the reduced levels of LC3-II, p62 and LAMP1 proteins in cardiac tissue with no major effect in quadriceps muscle (Figure 5A). On the other hand, Gys1 ASO#2, alone or in combination with ERT, led to the almost complete normalization of the p62, LC3-II and LAMP1 protein levels in quadriceps of Gaa^-/-^ mice (Figure 5A). In the heart, the Gys1 ASO contributed to a lesser extent to the reduction of the autophagic markers (Figure 5A). In addition, immunostaining of histological sections confirmed that ASO#2+ERT administration reduced LAMP1 protein levels in quadriceps muscle of Gaa^-/-^ mice (Figure 5A, 5B), suggesting that the combination therapy could rescue the autophagy pathway dysfunction in Pompe muscle after four months of treatment. The effect was more pronounced than in the Gys1 ASO#2-only group (Figure 5B).

**Figure 5.**
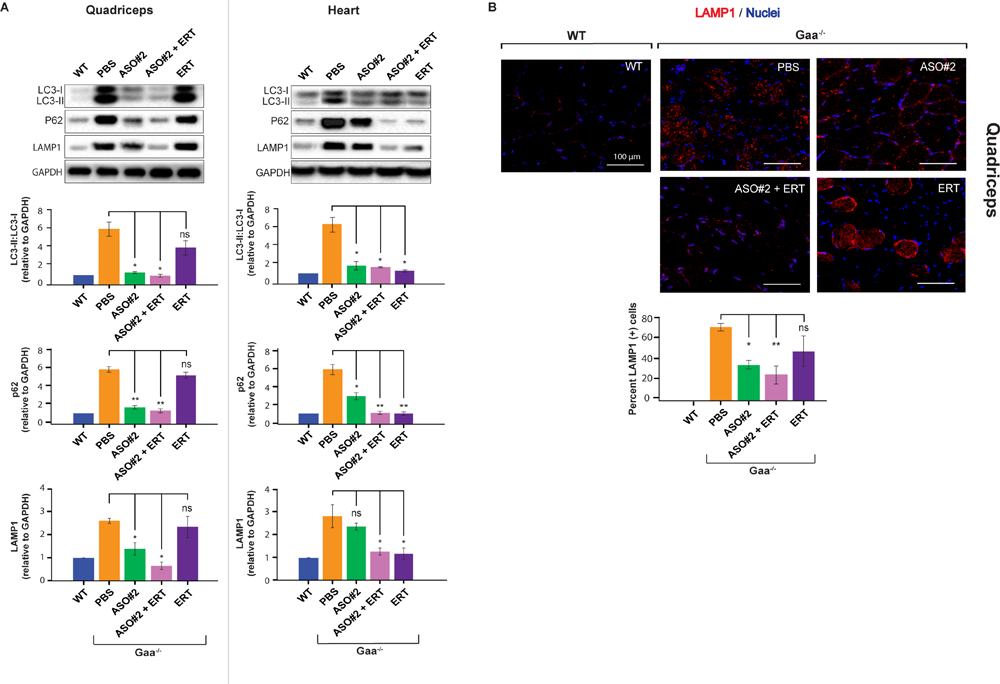
Normalization of autophagic flux in skeletal muscle of Pompe mice after treatment with Gys1 ASO, alone or in combination with ERT. **A.** Western blot analysis of LC3-II/LC3-I, p62, and LAMP1 proteins in quadriceps muscle and heart from mice in cohort 3. GAPDH was used as loading control. The graphs illustrate the quantification of Western blots from three independent experiments. WT: wild type. **B.** Representative images of LAMP1 immunostaining in histological sections of quadriceps muscle from Gaa^-/-^ mice treated with Gys1 ASO#2, alone or in combination with ERT. The graph shows the quantification of the stain as percentage of LAMP1-positive cells. LAMP1: lysosomal-associated membrane protein 1. Statistical analysis was performed using unpaired t-test. *: p<0.05, **: p < 0.01, ***: p < 0.005, ns: not significant.

**Figure 6:**
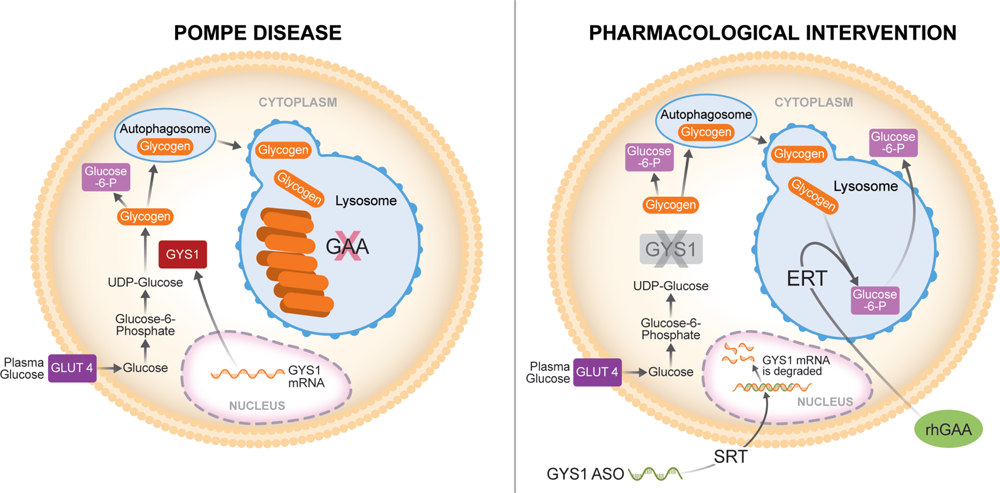
Model of pharmacological intervention in Pompe disease combining GYS1 ASO-based SRT and ERT. We propose an adjunct therapeutic approach for Pompe disease based on the reduction of the synthesis of new glycogen via GYS1 ASO-mediated SRT, combined with the ERT-mediated clearance of pre-existing glycogen that has accumulated in the lysosome because of GAA inactivation. Such combinatorial strategy would result in greater clinical benefit for Pompe patients, especially in skeletal muscle, where ERT alone is not remarkably effective. SRT: substrate reduction therapy; ERT: enzyme replacement therapy; GYS1: glycogen synthase 1; ASO: antisense oligonucleotide; GAA: glucosidase, alpha acid; rhGAA: recombinant human GAA.

In conclusion, our data indicate that the combination of Gys1 ASO and ERT might provide therapeutic benefit by targeting various aspects of Pompe pathogenesis, namely glycogen accumulation and autophagy defects.

### Systemic Gys1 knockdown does not mitigate Pompe pathology in the central nervous system

Pompe disease is a complex multi-systemic condition with growing evidence of central nervous system (CNS) involvement^35^. Indeed, we found remarkable abnormalities in the spinal cord of Gaa^-/-^ mice at 8 months of age (Supplementary Figure 3). The pathologic changes were visible in histological sections of the spinal cord, where Nissl staining (toluidine blue) revealed the presence of swollen cell bodies, activated glial cells, and centronucleated cells (Supplementary Figure 3). Systemically delivered Gys1 ASO#2 did not ameliorate the Pompe CNS phenotype of Gaa^-/-^ mice (Supplementary Figure 3), consistent with the inability of ASOs to cross the blood-brain barrier.

## DISCUSSION

Enzyme replacement therapy, currently the standard of care for patients with PD, has been shown to be beneficial, but its efficacy is hindered by some limitations, especially in treating the skeletal muscle manifestations of PD. Early administration of ERT improves cardiac hypertrophy and motor development, extending the chances of survival in patients with infantile-onset disease (IOPD)^6,36^. Patients with the late-onset form (LOPD) have early increase and stabilization of their pulmonary and motor function, however after reaching a plateau there is continued decline in their motor function such as the 6 Minute Walk Test^8,9^. Progressive muscle weakness thus continues to be a problem for long-term survivors of PD^5,6^. An explanation is provided by studies that have shown that the delivery of the therapeutic enzyme to skeletal muscle is insufficient to reduce the glycogen content and eliminate the autophagic buildup^6^. Recently, a study using a new adjunct therapy based on the enzyme and chaperone ATB200/AT222 combination showed enhanced efficacy over the currently approved ERT in Gaa^-/-^ mice. ATB200/AT222 was able to normalize muscle glycogen content and reduced the levels of lysosomal and autophagosomal markers in the muscle of Gaa^-/-^ mice to wild type levels^7,18^. Clinical trials of AT2221/ATB200 have been completed and this approved treatment now offers an alternative effective therapeutic option for Pompe disease patients.

Douillard-Guilloux et al. (2008) demonstrated that intramuscular injection of AAV-1 expressing shRNA targeted to Gys1 reduces glycogen accumulation in the gastrocnemius of newborn Gaa^-/-^ mice^19^. The same research group also analyzed the effect of a complete genetic ablation of glycogen synthesis in a new Gaa/Gys1 double knock-out (KO) mouse model^17^. The Gaa/Gys1 KO mouse exhibited a profound reduction in cardiac and skeletal muscle glycogen, a significant decrease in lysosomal swelling and autophagic buildup, and complete correction of cardiomegaly^17^. Muscle atrophy, observed in 11-month-old single Gaa KO mice, was less pronounced in the Gaa/Gys1 double KO mice, resulting in improved exercise capacity^17^. This data demonstrated that long-term suppression of muscle glycogen synthesis is well tolerated and leads to a significant improvement in Pompe pathology. However, no approved small molecule inhibitors of muscle GYS1 are currently clinically available, underscoring the challenges of finding specific compounds that can preserve the activity of the hepatic GYS2 enzyme.

ASO technology has emerged as a highly specific therapeutic option for a variety of neuromuscular diseases. ASOs that modulate exon splicing of the dystrophin (DMD) gene can reduce muscle degeneration in Duchenne’s muscular dystrophy^37–39,11,40,41^. An ASO that disrupts spliceosome action on exon 7 of the survival motor neuron 1 (SMN1) gene and restores the gene to functioning length has been approved for treatment of spinal muscular atrophy^42,43^. Recently, an ASO with RNase H1 mechanism has been approved by the FDA for amyotrophic lateral sclerosis (ALS) patients with SOD1 mutations^41,42^. Several studies have attempted to use ASOs to mitigate the disease outcomes by improving or restoring normal GAA synthesis^6^. Van der Wal et al. (2017) studied the common IVS132-13T>G GAA mutation and located sequences where an ASO could block splicing machinery and prevent exclusion of exon 2 in the final transcript, leading to increased GAA expression^44,45^. Goina et al. (2017) performed a similar exon 2 splicing modulation with an ASO and successfully reduced glycogen accumulation in patient-derived myotubes^46^. Other ASO studies have sought to reduce glycogen by interfering with GYS1 expression. Clayton et al. (2014) demonstrated that treating Pompe mice with a phosphorodiamidate morpholino oligonucleotide (PMO) designed to induce exon 6 skipping in the murine Gys1 mRNA resulted in the degradation of messenger RNA due to frame shift^16^. However, to facilitate PMO delivery to muscle via systemic administration, the PMO was conjugated with a cell penetrating peptide (GS-PPMO), and the development of PPMO drugs for human use is significantly challenged by their potential toxicity, as observed in primate studies^46^. This toxicity is likely due to the cationic nature of the peptide at the doses required for efficacious exon skipping in the primate musculature^16,47^. Benefiting from advancements in ASO technology, here we identified a potent ASO that safely knocks down mouse Gys1 expression by RNase H1 mechanism in mice. Using both prevention and reversal treatment modalities, we demonstrated the potential of the Gys1 ASO approach to serve as efficacious SRT in the Gaa^-/-^ mouse model of Pompe disease.

The Gaa^-/-^ Pompe mice start to accumulate glycogen in skeletal muscle as early as three weeks of age but develop a progressive defect in motor functions from six months of age. We have attempted to address the importance of timing in two separate cohorts, starting the ASO treatment at one-and three-month-old mice respectively. Taken together, our data indicate that long-term elimination of muscle glycogen synthesis leads to a significant reduction of glycogen accumulation and autophagic buildup, as well as a tendency towards functional improvement in severely affected older Pompe mice. Interestingly, we observed superior activity of Gys1 ASO in the skeletal muscle of Pompe mice compared to wild type, healthy mice (Supplementary Figure 1). We hypothesize this phenomenon being the result of increased permeability to ASO of the damaged muscle combined with facilitated escape of the ASO from defective endosomal compartments upon cellular uptake.

Despite the positive results obtained by knocking down Gys1 expression with ASO, there was still a residual amount of glycogen present in the skeletal muscle of older Gaa^-/-^ mice, and the muscle structure abnormality was not significantly improved upon treatment with ASO alone (Supplementary Table 2). This could be due to compensatory mechanisms, or very low levels of residual GYS1 enzymatic activity. To further improve the efficacy of our Gys1 ASO-based SRT approach both in the skeletal muscle and heart, we decided to combine the ASO treatment with ERT. Our data shows that the combination of Gys1 ASO with rhGAA results in enhanced reduction of glycogen content and reversal of the autophagy defect in the skeletal muscle of old Gaa^-/-^ mice. The molecular changes induced by the ASO and ERT treatment led to the preservation of motor function in the aging Pompe mice. Recently Ullman et al. (2024)^48^ recently reported their preclinical studies of MZ-101, a selective and noncompetitive inhibitor of GYS1, reduced glycogen accumulation, similar to that of ERT alone in Pompe mouse models. MZ-101 when utilized in combination with ERT however had an additive effect of normalizing glycogen concentrations in the muscles. These data from prior studies and our promising data suggests that substrate reduction therapy with GYS1 inhibition may be a promising therapeutic approach for Pompe disease. Recently, Maze Therapeutics announced positive phase 1 study results from a clinical trial of the small molecule GYS1 inhibitor MZE001 (ClinicalTrials.gov Identifier: NCT05249621). In the study the healthy volunteers demonstrated exposure-dependent reductions in blood mononuclear cell glycogen across dose levels 10 days after administration. It is also expected that GYS1 ASOs has the potential to be used as monotherapy or in combination with ERT in future clinical trials in Pompe patients. The results from the clinical trial further support the validity and safety of GYS1 inhibition as a potential therapeutic approach for Pompe disease.

## MATERIALS AND METHODS

### Identification of antisense oligonucleotides (ASOs) targeting mouse Gys1

All ASOs used in the studies described here were 16 nucleotides in length with a phosphorothioate (PS) backbone and three 2’-constrained ethyl (cEt)-modified nucleotides at both ends (3-10-3 gapmer configuration). A well-characterized control ASO which does not hybridize to any mouse mRNA sequence was included in the experiments. The oligonucleotides were synthesized and purified as previously described^49^. ASOs targeting murine Gys1 were designed and initially screened for activity in B16-F10 cells via electroporation. The most active ASOs were subsequently evaluated in wild type mice to identify ASOs that were safe and active in vivo. A dose-response confirmation experiment for the two lead ASOs, ASO#1 and ASO#2, was performed in wild type mice by subcutaneous injection twice a week for 5.5 weeks at 12.5, 25, 50, and 100 mg/kg total weekly doses (shown in Supplementary Figure 1A). All ASOs were formulated in phosphate buffered saline (PBS) and were administered to mice by subcutaneous injection. The mouse ASO screening studies were conducted under protocols approved by the Institutional Animal Care and Use Committees at Ionis Pharmaceuticals.

### Characterization of the Gaa^-/-^ animal model of Pompe disease

The Gaa^-/-^ Pompe mouse model was a gift of Dr. Nina Raben^50^. The mouse model is a constitutive homozygote knockout of the murine acid alpha-glucosidase (Gaa) gene. One-month-old Gaa^-/-^ mice, which show only the initial signs of tissue glycogen accumulation without severe myopathy, were used in prevention studies, whereas three-month-old Gaa^-/-^ mice, which show marked glycogen accumulation and motor function impairment, were used in reversal studies. All the intervention studies performed in either the Gaa^-/-^ mouse model or wild type C57BL/6 control mice were conducted in accordance with the Institutional Animal Care and Use Committee at the University of California, Irvine (UCI), under protocol IACUC AUP-19-075, and were consistent with Federal guidelines. The mice were housed at UCI animal facility that maintained a constant temperature (22°C) and controlled light and dark cycles (12:12). The overall condition of the mice was monitored daily throughout the experimental process to assess general health and mitigate any pain and suffering.

### Administration of ASOs to Gaa^-/-^ mice

Gaa^-/-^ or wild type control mice were dosed with ASOs once a week at 25 mg/kg via subcutaneous injection. For the early-treatment study (Cohort 1) described in Figure 2, one-month-old Gaa^-/-^ mice were randomly divided into groups for treatment with different Gys1 ASOs, control ASO, or vehicle (PBS), n=12-17 mice per group. For the reversal study (Cohort 2) described in Figure 3, three-month-old Gaa^-/-^ mice were randomly divided into three groups for treatment with Gys1 ASO#2 (n=15), control ASO (n=10), or vehicle (PBS) (n=10). The mice received a total of 16 weekly ASO doses (16 weeks duration) and were sacrificed 48 hours after the last ASO dose. Body weight changes, and the weight of liver, kidney, and spleen were recorded to exclude potential toxicity of the ASOs in the context of the long dosing paradigm in diseased Gaa^-/-^ mice.

### Adjunct treatment using Gys1 ASO in combination with ERT in Gaa^-/-^ mice

Four-month-old Gaa^-/-^ mice were dosed with Gys1 ASO#2 or control ASO by subcutaneous injection once a week at 25 mg/kg for six weeks. Starting from week 7 the mice additionally received 20 mg/kg recombinant human GAA (rhGAA) (Genzyme Corporation, Cambridge, MA; Cat.: NDC 58468-0160-1) via intravenous injection once every two weeks. The antihistaminic agent diphenhydramine hydrochloride was administered to the mice via intraperitoneal injection 15 minutes before receiving the human enzyme, starting from the second rhGAA administration. The weekly subcutaneous ASO administration continued throughout the ERT phase for a total of 16 doses. The mice were euthanized at 8 months of age, after having been treated with Gys1 ASO for 4 months, the last two of which overlapped with the ERT; n=5 per treatment group.

### Accelerating rotarod performance test

Muscle function was evaluated using a rotarod apparatus (Rotamex-5, Columbus Instruments, Columbus, OH). Briefly, mice were placed on a rotarod whose speed was set to progressively increase from 4 to 40 rotations per minute (rpm) in 300 seconds. The latency to fall from the rotating rod was recorded as a measure of muscle strength. Assessments were done on three consecutive days, three to five trials each day with at least two-minute rest intervals between the trials. The mean latency time to fall off the rotarod from three independent trials from the last two days was used for analysis.

### Biochemical analysis of serum from mice for toxicity studies

Blood samples were collected from mice via cardiac puncture under deep anesthesia using isoflurane inhalation. Immediately after collection, approximately 300-500 µL of whole blood were placed into serum separator tubes (Greiner Bio-One Minicollect tubes; Cat.: 450472). Samples were allowed to clot for at least 20 minutes at room temperature, and then centrifuged at 3,000 rpm for 15 minutes. Serum was transferred to fresh tubes and stored at −80°C before being used for clinical biochemistry evaluation at IDEXX Laboratories (IDEXX Laboratories Inc., Maine, USA). Parameters measured include alanine aminotransferase, aspartate aminotransferase, alkaline phosphatase, blood urea nitrogen, creatine kinase.

### Gene expression analysis using quantitative real-time PCR

Tissue samples were isolated from mice and snap frozen in liquid nitrogen. Frozen tissues were homogenized in TRIzol reagent (Thermo Scientific, Waltham, MA; Cat.: 15596026), and total RNA was extracted according to the manufacturer’s instructions. Real-time qPCR analysis was performed using the following reagents: mouse Gys1 (Taqman™ assay ID: Mm01962575_s1), mouse Gapdh (Taqman™ assay ID: Mm 99999915_g1, housekeeping gene). In the initial assessment of ASO activity in wild type mice (Supplementary Figure 1A) real-time qPCR analysis was performed using the following primer-probe sets: mouse Gys1 (forward primer: 5’-TGATGAAGAGAGCCATCTTTGC-3’; reverse primer: 5’-AGGAGTCGTCCAGCATGTTGT-3’; probe: 5’-ACTCAGCGGCAGTCTTTCCCACCA-3’), mouse cyclophilin A (forward primer: 5’-TCGCCGCTTGCTGCA-3’; reverse primer: 5’-ATCGGCCGTGATGTCGA-3’; probe: 5’-CCATGGTCAACCCCACCGTGTTC-3’). Reactions were conducted in triplicate on a QuantStudio7 real-time PCR system (Thermo Fisher Scientific, Waltham, MA), following the manufacturer’s instructions.

### Biochemical measurement of tissue glycogen content

Mouse tissues were collected, snap frozen in liquid nitrogen upon collection, and stored at −80°C. Approximately 10 mg of frozen quadriceps muscle were rinsed in 300 μL of ice-cold PBS, then homogenized in 200 μL of double-distilled water using a Wheaton dounce tissue grinder (DWK Life Sciences, Rockwood, TN) as follows: 10-20 passes on loose pestle followed by 10-20 passes on tight pestle, keeping the tissue grinder on ice. The tissue lysates were then heated at 100°C for 10 minutes, then centrifuged at 15,000 g at 4°C for 10 minutes. Total protein concentration in the supernatant was determined using the Micro BCA protein assay kit (Thermo Scientific, Waltham, MA; Cat.: 23235). In the same supernatant, glycogen concentration was measured using a fluorometric enzymatic assay (Sigma-Aldrich, St. Louis, MO; Cat.: MAK016), according to the manufacturer’s instructions. The assay produces a fluorometric (λex = 535/λem = 587 nm) product proportional to the glycogen present in the sample. The fluorescent signal was analyzed using the BioTek Gen5 software (Agilent Technologies, La Jolla, CA), and normalized by the total protein content of the sample.

### Western blot analysis of autophagy markers

Approximately 30 mg of frozen tissue was homogenized in 500 μL of RIPA buffer (Sigma-Aldrich, St. Louis, MO; Cat.: R0278) supplemented with a proteinase inhibitor cocktail (Sigma-Aldrich, St. Louis, MO; Cat.: P8340) using a handheld electric homogenizer (Qsonica, Newtown, CT; Cat.: XL-2000). The tissue homogenate was then agitated on a Benchmark Roto-Mini Plus (Benchmark Scientific, Sayreville, NJ) for 2 hours at 4°C. Lastly, the tissue lysate was centrifuged at 20,000 g at 4°C for 30 minutes, and the supernatant was collected for analysis. The sample protein concentration was measured using the Micro BCA protein assay kit (Thermo Fisher Scientific, Waltham, MA; Cat.: 23235). 20 μg of total protein lysate was resolved by electrophoresis on Bis-Tris 4-12% NuPAGE polyacrylamide gels using the Novex Mini Cell system (Invitrogen/Thermo Fisher Scientific, Waltham, MA). The proteins were then transferred to a PVDF membrane, which was later blocked with 1% bovine serum albumin (BSA) in Tris-buffered saline containing 0.1% Tween 20 (TBS-T). The primary antibodies used in our studies were: anti-GYS1 (Abcam, Waltham, MA; Cat.: ab 40810; 1:5,000 dilution), anti-LC3B-I/II (Abcam, Waltham, MA; Cat.: ab192890 ; 1:5,000 dilution), anti-p62 (Abcam, Waltham, MA; Cat.: ab56416; 1:20,000 dilution), anti-LAMP1 (Abcam, Waltham, MA; Cat.: ab27170; 1:2,000 dilution), anti-GAPDH (Abcam, Waltham, MA; Cat.: ab18160; 1:10,000 dilution; served as loading control), anti-β-ACTIN (Abcam, Waltham, MA; Cat.: ab8227; 1:5,000 dilution; served as alternative loading control). Horseradish peroxidase (HRP)-linked goat anti-mouse or goat anti-rabbit secondary antibodies were used (1:5,000 dilution). Densitometric quantification was performed using ImageJ software (National Institutes of Health, Bethesda, MD).

### Periodic acid-Schiff Histological analysis

Periodic acid-Schiff (PAS) staining was performed to detect glycogen in histological tissue sections. Briefly, the quadriceps muscle was dissected and fixed in a 4% neutral buffered formalin solution at room temperature for 20 hours. The fixed tissue was then rinsed in PBS and transferred to a 70% ethanol solution before being embedded in paraffin blocks and sectioned. The PAS staining was performed on 8μm-thick sections by the Pathology Department at UCI Medical Center according to protocols described previously^49^. Images of the histological slices were captured using a Keyence BZ-810 microscope (Keyence, Itasca, IL). In PAS-stained tissue sections, glycogen appears fuchsia-colored under light microscopy. For quantification of the PAS staining, the histological sections of quadriceps muscle from 4-5 animals per treatment group were analyzed, and a semi-quantitative score from 1 (no staining) to 4 (extensive staining) was assigned to each section. The scoring was performed independently by two different technicians to ensure accurate assessment. To quantify the number of myofibers with centralized nuclei, a minimum of 300 myofibers per animal were manually evaluated in the PAS-stained sections of quadriceps muscle (n=4-5 mice per each group). The data is presented as the percentage of myofibers with a centralized nucleus.

### Immunohistochemistry (IHC) studies

For lysosomal analysis, the expression of lysosomal associated membrane protein 1 (LAMP1) was assessed using IHC in tissue sections of quadriceps muscle embedded in frozen section medium (Fisher Scientific, Hampton, NH; Cat.: 22-046-511), as described previously^51^. Briefly, 8µm-thick cryosections (10 sections per each animal, 5 animals per treatment group) were fixed in 4% paraformaldehyde (PFA), blocked in donkey serum, and stained with an anti-LAMP1 primary antibody (Abcam, Waltham, MA; Cat.: ab24170; 1:500 dilution) at 4°C overnight. The sections were then washed three times with PBS and incubated with fluorescein-conjugated secondary antibody (Thermo Fisher Scientific, Waltham, MA; Cat.: A10042; 1:500 dilution) for 1 hour at room temperature, then washed and mounted with DAPI-containing mounting media (Vectashield, Vector Laboratories, Newark, CA; Cat.: H-1200-10). The slides were imaged on a Zeiss LSM 900 confocal microscope (Carl Zeiss Microscopy, White Plains, NY). For the quantification of the LAMP1 staining, the percentage of LAMP1-positive myofibers was calculated and compared across treatment groups.

Nissl (toluidine blue) staining was used to study the morphology and pathology of neural tissue. The spinal cord was harvested from mice, fixed in 4% paraformaldehyde (PFA) for 24 hours, and then incubated in a solution containing 30% sucrose in PBS overnight, embedded in optimal cutting temperature compound (OCT compound), and cryo-sectioned. Subsequently, the 10-micron cryosections were stained with 1% toluidine blue containing 1% borate solution as previously described^52,53^. Briefly, the slices were placed in Toluidine blue solution for 30 seconds and rinsed in triple distilled water. Stained slices were examined using light microscopy.

### Statistical analysis

Statistical analysis was performed using GraphPad Prism software (GraphPad Software, Boston, MA), or RStudio software (RStudio, Boston, MA) in the case of Figure 1B. Unpaired t-test, one-way ANOVA, or two-way ANOVA were used, as specified in the respective figure legends. p-values lower than 0.05 were considered statistically significant; *: p<0.05, **: p<0.01, ***: p < 0.005, and ****: p<0.0001, ns= not significant. Error bars indicate the standard error of the mean (SEM).

#### List of abbreviations

ASO: antisense oligonucleotide
GYS1: glycogen synthase 1
GAA: glucosidase, alpha acid
rhGAA: recombinant human GAA
SRT: substrate reduction therapy
ERT: enzyme replacement therapy
PD: Pompe disease

## ACKNOWLEDGEMENTS

We thank the NIH/NIAMS R21 AR080972-01) (PI Kimonis), Helen Walker Award, AMDA (Acid Maltase Deficiency Association), and the Council on Research, Computing, and Libraries (CORCL), UC Irvine, for funding this study. We thank Jillian Vu, Marvan Youssef and the UCI ULAR staff for their assistance with the animal experiments. We also thank Tracy Reigle and Raul Alonzo (Ionis Pharmaceuticals) for assistance with the preparation of figures.

---

**Supplementary Figure 1:**
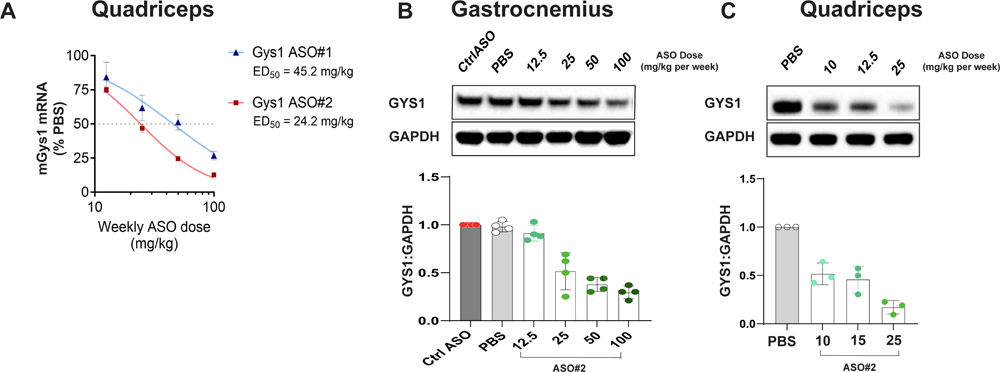
Gys1 ASO#1 and ASO#2 knock down mouse Gys1 mRNA and protein in wild type mice. **A.** Knockdown of mouse Gys1 mRNA measured by qPCR in quadriceps muscle of wild type mice treated with ASO#1 and ASO#2 by subcutaneous injection at the indicated total weekly doses; n=4 per group. **B.** Western blot analysis of GYS1 protein in gastrocnemius muscle of wild type mice treated via subcutaneous injection with ASO#2 for 6 weeks at the indicated total weekly doses; n=4 per group. The graph shows the quantification of GYS1 protein expression relative to GAPDH (loading control) from three independent experiments. The GYS1 expression level after treatment with control ASO (Ctrl ASO) at 100 mg/kg total weekly dose was used as reference and was therefore set to 1. **C.** Western blot analysis of GYS1 protein in quadriceps muscle of Gaa^-/-^ mice treated via subcutaneous injection with ASO#2 for 6 weeks at the indicated weekly doses; n=3 per group. The PBS control group was used as reference to determine normal GYS1 levels and was therefore set to 1. Representative blots from 3 sets of experiments are shown.

**Supplementary Figure 2.**
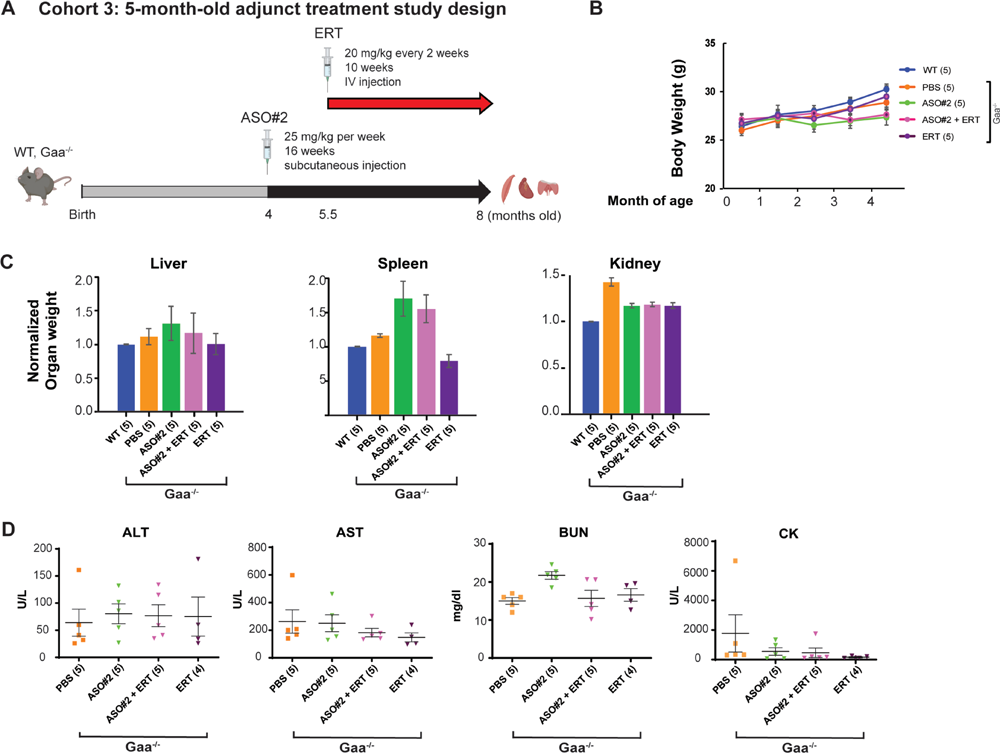
Schematic design of the treatment strategy and toxicity studies using Gys1 ASO alone or in combination with ERT in Gaa-/-mice. Gys1 ASO and ERT are well tolerated in Gaa^-/-^ mice. **A.** Schematic illustration of the study used to assess the safety of the Gys1 ASOs in vivo. **B.** Change in body weight of wild type and Gaa^-/-^ mice dosed with either PBS, Gys1 ASO#2 alone, or Gys1 ASO#2 in combination with ERT. The mice were dosed from 4 to 8 months of age, and their body weights were recorded once every four weeks during the four-month study. **C.** Terminal organ weight to body weight ratio normalized to wild type control group (set to 1). All the treatments were well tolerated since the ratios are within two-fold of wild type. **D.** Serum toxicology analysis performed at the end of the study shows no statistically significant change in the treatment groups compared to control Gaa^-/-^ mice dosed with PBS. The number of samples in each group is listed in parentheses on the x-axis. ALT: alanine transaminase (values above 100 U/L are considered a sign of liver toxicity); AST: aspartate transaminase (values above 250-300 U/L are considered a sign of liver toxicity); BUN: blood urea nitrogen (values above 30 mmol/L are considered a sign of kidney toxicity); CK: creatine kinase (values above 1,500 U/L are considered a sign of muscle toxicity).

**Supplementary Figure 3.**
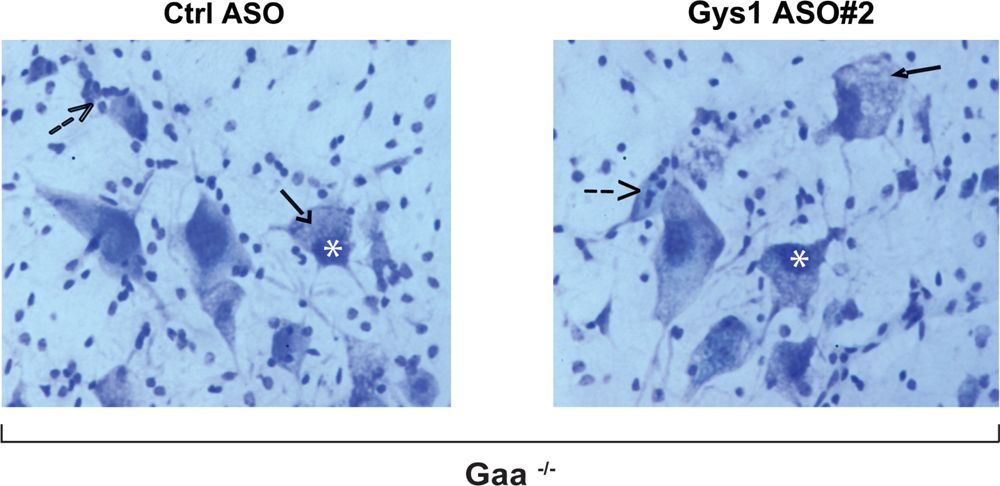
Histological analysis of spinal cord in 8-month-old Gaa^-/-^ mice with systemically delivered Gys1 ASO. No effect of systemically delivered Gys1 ASO in CNS. Nissl (toluidine blue) staining of histological sections of spinal cord from 8-months-old Gaa^-/-^ mice dosed by subcutaneous injection with 25 mg/kg of either control ASO or Gys1 ASO#2 once a week for 16 weeks (16 total doses, 3.5 months of treatment, Cohort 2). Solid arrows indicate swollen cell bodies in the cervical ventral horn; dashed arrows indicate activated glial cells; asterisks indicate eccentric nuclei.

**Supplementary Table 1.** Summary of percent reduction over control ASO or PBS of Gys1 mRNA, glycogen content, and GYS1 protein in Gaa^-/-^ mice treated with ASO, ASO+ERT, or ERT alone. Mice from three separate cohorts were dosed with the indicated treatment, starting at 1, 3, or 4 months of age, and ending at 4.5, 6.5, or 8 months of age, respectively. The data is reported as percent reduction over control ASO or PBS of Gys1 mRNA, glycogen content, and GYS1 protein in Gaa^-/-^ mice treated with ASO, SO+ERT, or ERT alone

|  |  | % reduction |  |  |  |  |  |
| --- | --- | --- | --- | --- | --- | --- | --- |
|  |  | Gys1 mRNA<br>(±95% C.I.) | GYS1 protein |  |  | Glycogen content<br>(±95% C.I.) |  |
| Mouse cohort | Treatment | Quadriceps | Quadriceps | Diaphragm | Heart | Quadriceps | Heart |
| 1-month-old Gaa<sup>-/-</sup><br>(vs. Control ASO) | ASO#1<br>(n=12) | 83.64<br>(77.40 - 88.16) | 98 | 85 | 90 | 38.06<br>(14.50-61.07) | N.M. |
|  | ASO#2<br>(n=12) | 88.58<br>(83.61 - 92.06) | 100 | 98 | 100 | 57.05<br>(35.57-71.36) | N.M. |
| 3-month-old Gaa<sup>-/-</sup><br>(vs. Control ASO) | ASO#2<br>(n=10) | 64.71<br>(57.00 - 71.05) | 100 | 80 | 70 | 7.97<br>(5.53-28.45) | N.M. |
| 4-month-old Gaa<sup>-/-</sup><br>(vs. PBS) | ASO#2<br>(n=5) | N.M. | 100 | 90 | 100 | 55.22<br>(20.29-74.84) | 14.03<br>(28.58-47.22) |
|  | ASO#2+ERT<br>(n=5) | N.M. | 100 | 85 | 90 | 83.48<br>(53.80-94.10) | 65.82<br>(5.88-87.59) |
|  | ERT<br>(n=5) | N.M. | 0 | 0 | 0 | 15.78<br>(11.61-31.49) | 74.93<br>(18.77-92.26) |

N.M.: not measured; C.I: confidence interval.

**Supplementary Table 2.**
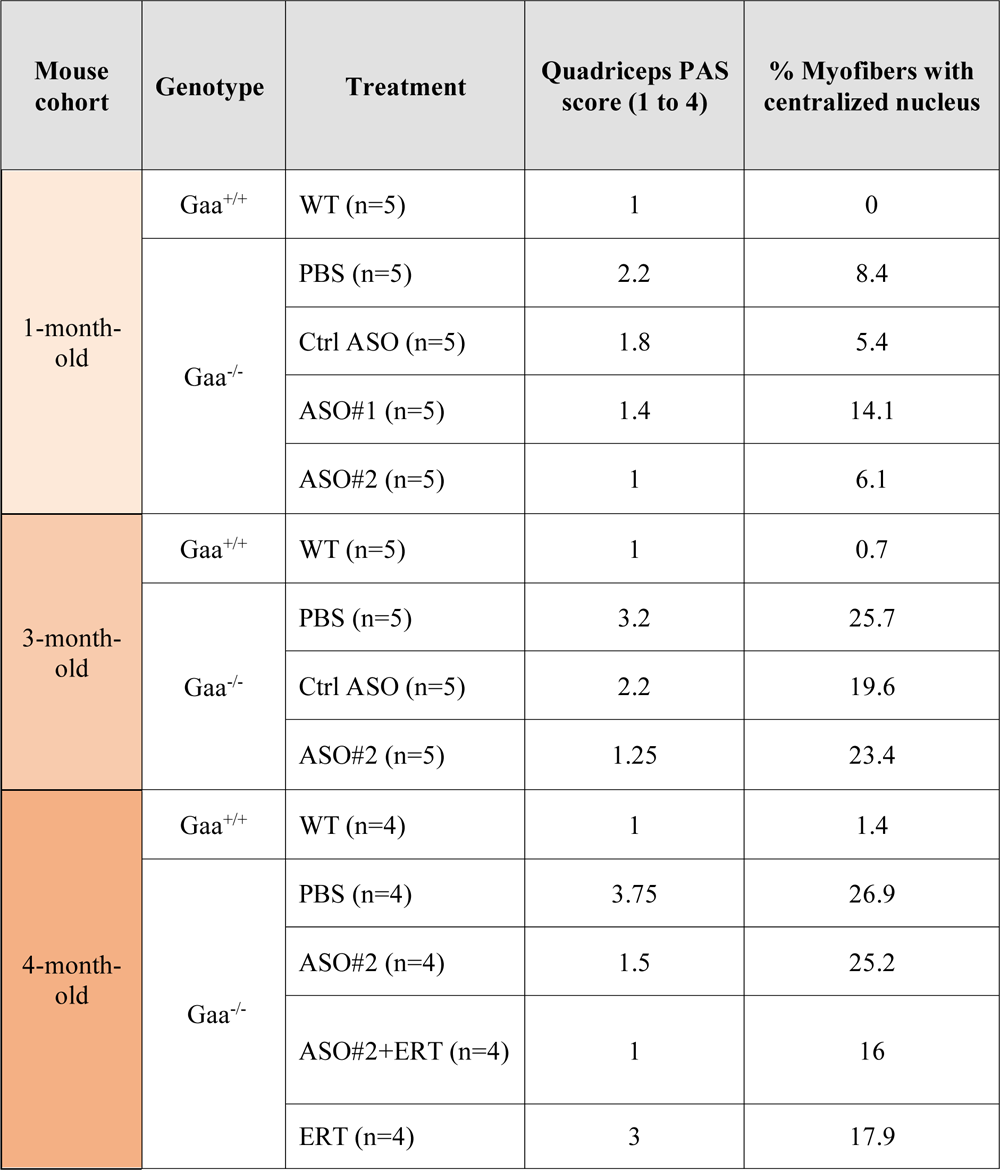
Summary of histological analyses. Semi-quantitative analysis of PAS staining, and percentage count of myofibers with a centralized nucleus.

